# *LorMe*: a streamlined and interoperable R framework for end-to-end microbiome analysis

**DOI:** 10.1101/2025.11.26.690896

**Authors:** Ningqi Wang, Yaozhong Zhang, Xinrun Yang, Yang Gao, Hang Xia, Yangchun Xu, Qirong Shen, Alexandre Jousset, Gaofei Jiang, Zhong Wei

## Abstract

Microbiome research is rapidly evolving accompanied by increasingly diverse analytical strategies and visualization formats. However, existing R-based tools remain fragmented, differ in data structures, and often require extensive package-specific pre-processing. This increases command-line complexity and limits both interoperability and reproducibility across packages. Here, we present *LorMe* (Lightweight One-line Resolving Microbial Ecology), an R package that provides a unified, interoperable, and user-friendly framework for microbiome data analysis. *LorMe* incorporates a standardized S4 object system with fully bidirectional compatibility with *phyloseq* and *microeco*, which enables seamless data exchange across major microbial analysis packages. A global configuration system ensures consistent parameter control and visualization standards, and a modular architecture supports both complete end-to-end workflows and flexible execution of individual analytical components. Based on these, *LorMe* further offers an one-command pipeline that conducts the full spectrum of microbial community analyses, including alpha and beta diversity, differential abundance testing, co-occurrence network inference, and meta-network construction, while archiving all intermediate objects and source data to ensure complete reproducibility. Demonstrations using sample datasets show that *LorMe* performs comprehensive analyses within minute, produces publication-ready outputs, and maintains methodological transparency. Application to a real rhizosphere dataset revealed biologically coherent patterns across diversity metrics, differential taxa, and network modules, highlighting the capacity of *LorMe* to support robust ecological interpretation. Collectively, *LorMe* provides a lightweight, extensible, and flexible solution that reduces technical barriers and enhances reproducibility in microbiome research. *LorMe* package is freely accessible from both CRAN (https://cran.r-project.org/web/packages/LorMe) and github (https://github.com/wangnq111/LorMe)

## Introduction

Microbiome research has expanded rapidly across ecology, agriculture, medicine, and environmental sciences, yet its analytical workflows remain technically demanding and procedurally complex. Modern studies rely on high-throughput amplicon or metagenomic sequencing and typically involve several core analytical modules, including community profiling, differential abundance testing, and ecological network construction [1]. Each module offers multiple methodological options with distinct statistical assumptions and appropriate application scenarios [2]. For instance, differential abundance analysis alone includes more than ten commonly used approaches [3]. Beyond methodological diversity, microbiome pipelines generally require extensive data preprocessing and numerous parameter settings. Such complex workflows create high technical barriers for beginners and complicate standardization across studies [4]. As datasets grow larger and experimental designs become more complex, method heterogeneity and workflow configuration burdens increasingly hinder the selection of appropriate approaches and compromise reproducibility.

R remains the most flexible and widely adopted environment for microbiome analysis [5, 6], yet methodological challenges persist in two interconnected dimensions. First, despite numerous R packages supporting community profiling, differential abundance testing, and ecological network inference, differences in underlying design philosophies fragment the computational ecosystem. Widely used tools such as *phyloseq* [7], *microeco* [8], and MicrobiotaProcess [9] provide powerful functionalities, but their internal data structures are largely incompatible, forcing repeated object conversion and output reconciliation when integrating complementary methods. This structural heterogeneity also requires manual harmonization of visualization parameters and analytical settings. Second, parameter management and workflow assembly remain decentralized. Cloud-based platforms such as CFViSA [10], OmicStudio [11] partially address this challenge with preset or one-step pipelines, but implementing such designs directly in R functions would require embedding numerous configurable parameters into single functions, producing overly complex architectures and increasing the burden for both developers and users. Consequently, analyses often rely on user-assembled pipelines, and output formats remain inconsistent across tools. Together, these limitations reduce interoperability, complicate cross-method validation, and hinder reproducible research. These challenges highlight the need for a framework that unifies data representation, stabilizes interfaces across established packages, and supports both end-to-end and modular analyses while reducing technical burdens.

To address these limitations, we developed *LorMe* (Lightweight One-Line Resolving Microbial Ecology), a lightweight and interoperable R framework that unifies microbiome data representation and streamlines analytical workflows. *LorMe* introduces a standardized object-encapsulation system supporting two construction modes: direct integration of experimental design, multi-level taxonomic profiles, and phylogenetic information, and seamless conversion from *phyloseq* or *microeco* objects. This bidirectional compatibility allows *LorMe* to bridge existing R ecosystems without duplicating functionality. A global configuration system centralizes analytical and visualization parameters, while archiving of intermediate objects tracks each computational step. *LorMe* implements an integrated end-to-end pipeline performing community profiling, differential abundance testing, and ecological network analyses through a single command, while individual components can also be executed independently for flexible exploration. By combining lightweight design with comprehensive analytical coverage, *LorMe* reduces configuration burden, enhances reproducibility, and provides a unified framework that enables researchers to focus on biological interpretation rather than technical workflow assembly.

## 2. Materials and Methods

### 2.1 Architecture of *LorMe* object

The *LorMe* object is implemented using an S4-based framework. A specialized class definition is added so that all components can be accessed through both “$” and “@”, providing flexibility for users with different coding preferences. A *LorMe* object contains five core elements, including a metafile describing sample and treatment information, relative and absolute abundance tables for all taxonomic ranks from domain to species, taxonomy tables corresponding to these ranks, an optional phylogenetic tree, and a unified analytical configuration component.

During object construction, the function ‘tax_summary’ automatically generates abundance matrices across multiple taxonomic levels by hierarchically aggregating the lowest-level identifiers such as ASVs, OTUs, or metagenomic annotations. Both relative and absolute abundance tables are generated simultaneously, and corresponding taxonomic annotations are harmonized across taxonomic ranks. After configuration via ‘object_config’, analysis-related settings, including experimental design, color palettes, factor ordering, faceting rules, and paired-sample relationships, are stored within the object. All operations are executed through modular S4 methods to ensure internal consistency, controlled data access, and minimal dependence on the users’ environment. This structure provides a reproducible and stable foundation for all downstream analyses in *LorMe*.

### 2.2 Ecosystem interoperability

To ensure compatibility with established microbial ecology workflows, *LorMe* incorporates a bidirectional interoperability system that allows seamless conversion between *LorMe* objects and structures from widely used packages, including *phyloseq* and *microeco*. Four conversion utilities are provided, enabling the construction of *LorMe* objects from *phyloseq* or *microeco* formats as well as conversion of *LorMe* objects into these formats. All essential data layers, including abundance tables, taxonomy annotations, and metadata, are preserved during conversion. Because many microbial analysis tools (such as MicrobiotaProcess, ampvis2 [12], and vegan-based [13] ordination workflows) accept *phyloseq* objects as input, this interoperability layer allows *LorMe* to function either as a standalone analytical framework or as a complementary module within the broader microbiome R ecosystem.

### 2.3 Global configuration system

To simplify workflow control and minimize user workload, *LorMe* employs a unified global configuration system. The main analytical pipeline, implemented through *LorMe*_pipeline, requires only a *LorMe* object and a selected set of analytical steps, which default to executing all modules. Parameters for individual analyses are stored in a centralized global options structure with predefined default values. Users may modify these settings through the *LorMe*_options function, and *LorMe*_defaults restores the configuration to its initial state. All settings are bound to the package environment, ensuring reproducibility and preventing accidental overwriting by external variables. This configuration system standardizes visualization and reporting across modules while supporting rapid exploratory analysis and reproducible workflows

### 2.4 Modular system design

*LorMe* is organized into five major functional modules that share a unified architecture yet operate independently when needed (Figure 1A). The modular design allows users to execute complete workflows or invoke individual analytical components. All visualizations inherit the unified stylistic configuration stored in the *LorMe* object, ensuring consistency and reproducibility. These modules collectively form the *LorMe* microbial analysis pipeline.

**Figure 1.**
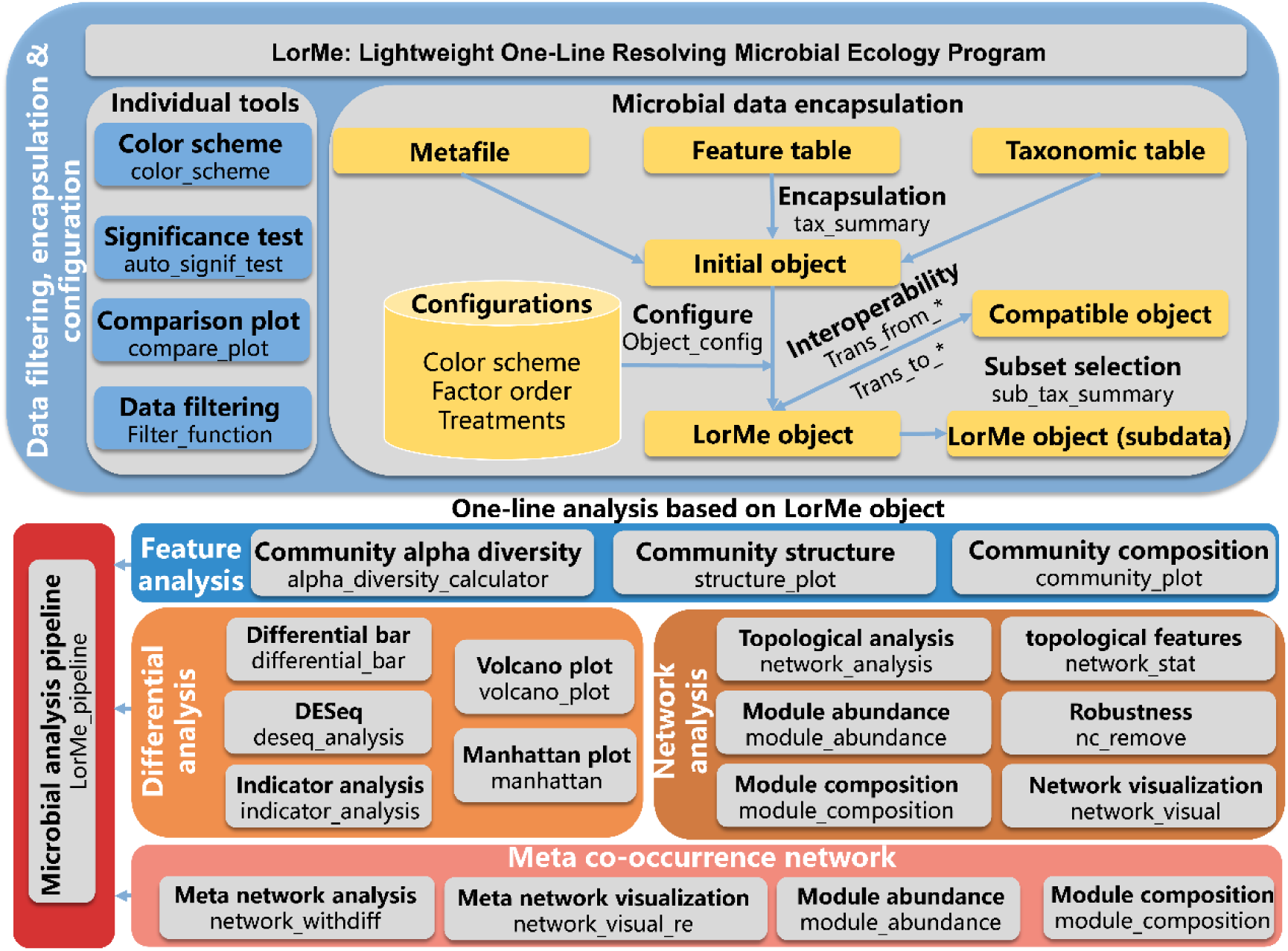
Overview of the *LorMe* package. The *LorMe* framework consists of five functional modules integrated into a single analytical pipeline. Colors represent different modules, bold labels denote analytical tasks, and regular text indicates corresponding *LorMe* functions.

#### 2.4.1 Data processing, encapsulation, and configuration

This module consists of two subcomponents. The first provides independent utilities for data filtering, color palette generation, significance testing, and simple multi-group visualizations. The second handles data encapsulation and configuration, including *LorMe* object construction, configuration management, subset extraction, and interoperability with external microbiome analysis packages. These tools ensure structural consistency of the data while supporting flexible preprocessing across diverse datasets.

#### 2.4.2 Community feature analyses

*LorMe* provides a comprehensive suite for quantifying microbial community characteristics, including α-diversity, β-diversity, and community composition. α-diversity metrics such as observed richness, Chao, ACE, evenness, Simpson, and Shannon indices are computed using vegan [13]. β-diversity and community structure are assessed using Bray–Curtis distances, followed by ordination methods including principal component analysis, principal coordinates analysis, and non-metric multidimensional scaling. statistics of permutational multivariate analysis of variance are provided to evaluate treatment-level differences. Community composition can be analyzed at all taxonomic ranks stored in the *LorMe* object, and visualized through bar plots, box plots, and alluvial diagrams. All analyses rely on the harmonized abundance matrices to maintain consistency across taxonomic resolutions.

#### 2.4.3 Differential abundance analyses

This module integrates several statistical approaches for differential analysis. Pairwise group comparisons are conducted using Wilcoxon tests with p-value adjustment by the Benjamini–Hochberg method, supported by built-in visualization tools. Differential abundance analysis for sequencing data is implemented using DESeq2 [14]. Indicator species analysis is performed through the indicspecies package [15]. Visualizations, including volcano plots and Manhattan-style representations, are provided to facilitate interpretation of results.

#### 2.4.4 Co-occurrence network construction and topological Analyses

*LorMe* contains a dedicated ecological network analysis system comprising six major functions. Co-occurrence networks are constructed using the Hmisc package to improve computational efficiency. Benjamini–Hochberg correction is applied to pairwise correlation tests [16]. Network modularization is computed using the fast-greedy algorithm via igraph. Additional functionalities include estimation of topological parameters, module-level abundance summarization, module composition analysis, robustness assessment, and network visualization. All computations are performed using the standardized abundance matrices, ensuring consistent taxonomic resolution throughout network analyses. The resulting networks can be visualized within *LorMe* or exported to external platforms such as Gephi and Cytoscape [17].

#### 2.4.5 Meta-Network Analyses

To integrate network structure and differential abundance results, *LorMe* provides a meta-network analysis module that incorporates global co-occurrence networks with taxonomic differences. This system facilitates the identification of treatment-associated changes in microbial interaction patterns and supports comparative visualizations such as overlaid networks and module-level differential summaries [18]. Given the exploratory nature of network-based interpretation, the module supports customized re-visualization according to user needs.

### 2.5 Implementation and software dependencies

*LorMe* is constructure in R (V4.4.1) using a streamlined dependency architecture designed to reduce installation burden and enhance reproducibility across computational environments. All the code and constructing principles followed standards in the comprehensive R archive network (CRAN). The package includes 21 core dependencies and 12 optional packages. This lightweight design minimizes version conflicts, shortens installation time, and ensures stable operation on both personal computers and high-performance computing environments. All functions are built using efficiency-oriented S4 methods, and no system-level libraries or external compilers are required. *LorMe* supports Linux, macOS, and Windows systems.

## 3. Results

### 3.1 *LorMe* workflow

*LorMe* provides a streamlined workflow that spans data preparation, object encapsulation, statistical analysis, and visualization, supporting the entire process of microbiome analysis (Figure 2). The framework consists of three major steps:

1. Users load the required input data. *LorMe* accepts three essential components: a feature table, a taxonomy table, and metadata describing the experimental design. A phylogenetic tree can be optionally included depending on analytical need.
2. The encapsulation and configuration step constructs the *LorMe* object. *LorMe* provides two routes for generating the preconfigured object. The direct-construction route derives hierarchical taxonomic profiles from domain to species on the basis of the feature and taxonomy tables and then integrates the metadata to generate the preconfigured object. Alternatively, *LorMe* allows users to convert an existing *phyloseq* or *microeco* object directly into a preconfigured *LorMe* object through built-in bidirectional conversion functions. After the preconfigured object is generated, analytical settings such as experimental design, color schemes, factor order, and visualization themes are applied to produce the final *LorMe* object. This configuration process ensures consistent analytical styles and visualization standards throughout downstream analyses.
3. The microbial analysis pipeline is executed using the *LorMe* object. The standardized pipeline integrates modular design with comprehensive analytical coverage, implementing alpha-diversity estimation, beta-diversity and community structure analysis, differential abundance testing, and co-occurrence network construction. These wrappers perform internal method selection, coordinate required parameters, and automatically format results, all under the global configuration system embedded in the package environment. Default parameters are initialized for immediate use, but users can freely adjust them to refine the final analytical settings. This design allows users to obtain publication-ready results with minimal scripting effort while preserving analytical flexibility.

**Figure 2.**
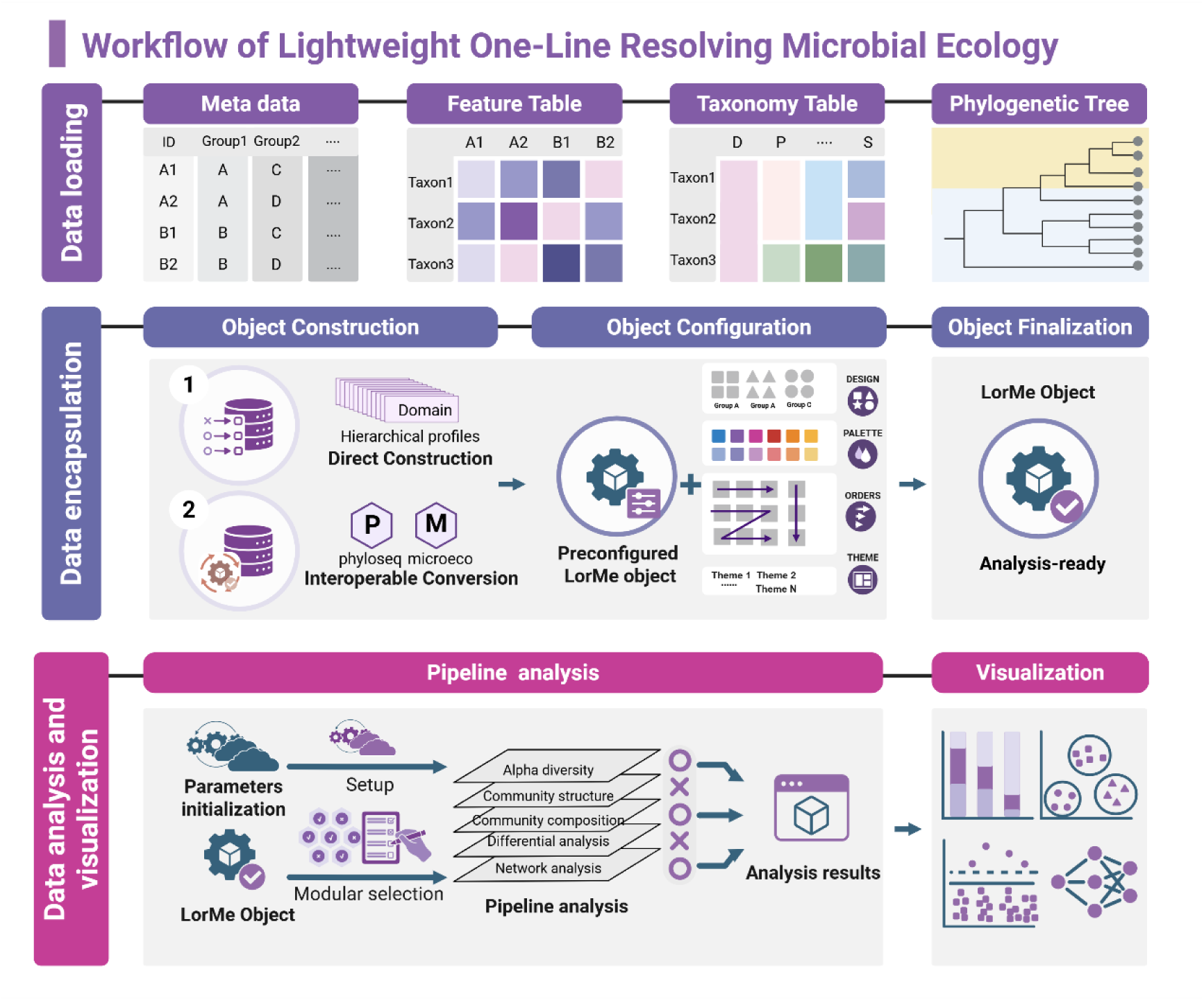
Overview of the analytical workflow implemented in *LorMe*. The workflow consists of three major steps: (1) Data loading. Metadata, feature tables, and taxonomy tables are required, whereas a phylogenetic tree is optional. (2) Data encapsulation. The loaded data are organized into hierarchical profiles and encapsulated into a preconfigured *LorMe* object, which serves as the direct construction method. Alternatively, existing *phyloseq* or micreco objects can be converted into the preconfigured *LorMe* objects. After construction, an analysis configuration is assigned, generating an analysis-ready *LorMe* object. (3) Data analysis and visualization. *LorMe* embedded a global parameter system in the package environment, where all analysis parameters are pre-initialized. After users customize the parameter settings, the *LorMe* object is processed through the analysis pipeline, with optional modules available for selection. The analytical results are summarized and can be directly visualized within *LorMe* or exported as source data for compatible downstream analyses and visualizations.

After the analysis is completed, all statistics, source data, visualizations, and intermediate results are archived within the output object to support downstream operations and external compatibility (see Supplemental Table 1 for a detailed output list). For example, users may refine plots using ggplot2, regenerate figures from extracted data, or export adjacency matrices and node tables for visualization in Gephi or Cytoscape. Overall, *LorMe* provides a lightweight yet comprehensive workflow that substantially reduces coding requirements while maintaining methodological rigor and the exploratory flexibility required for microbial ecological research.

### 3.1 Pipeline Demonstration

To illustrate the simplicity of *LorMe*, we demonstrate its one-command pipeline, which performs the complete microbiome workflow using a single command while maintaining full reproducibility and publication-grade output quality (Figure 3).

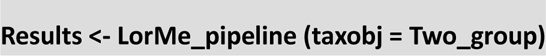

**Figure 3.**
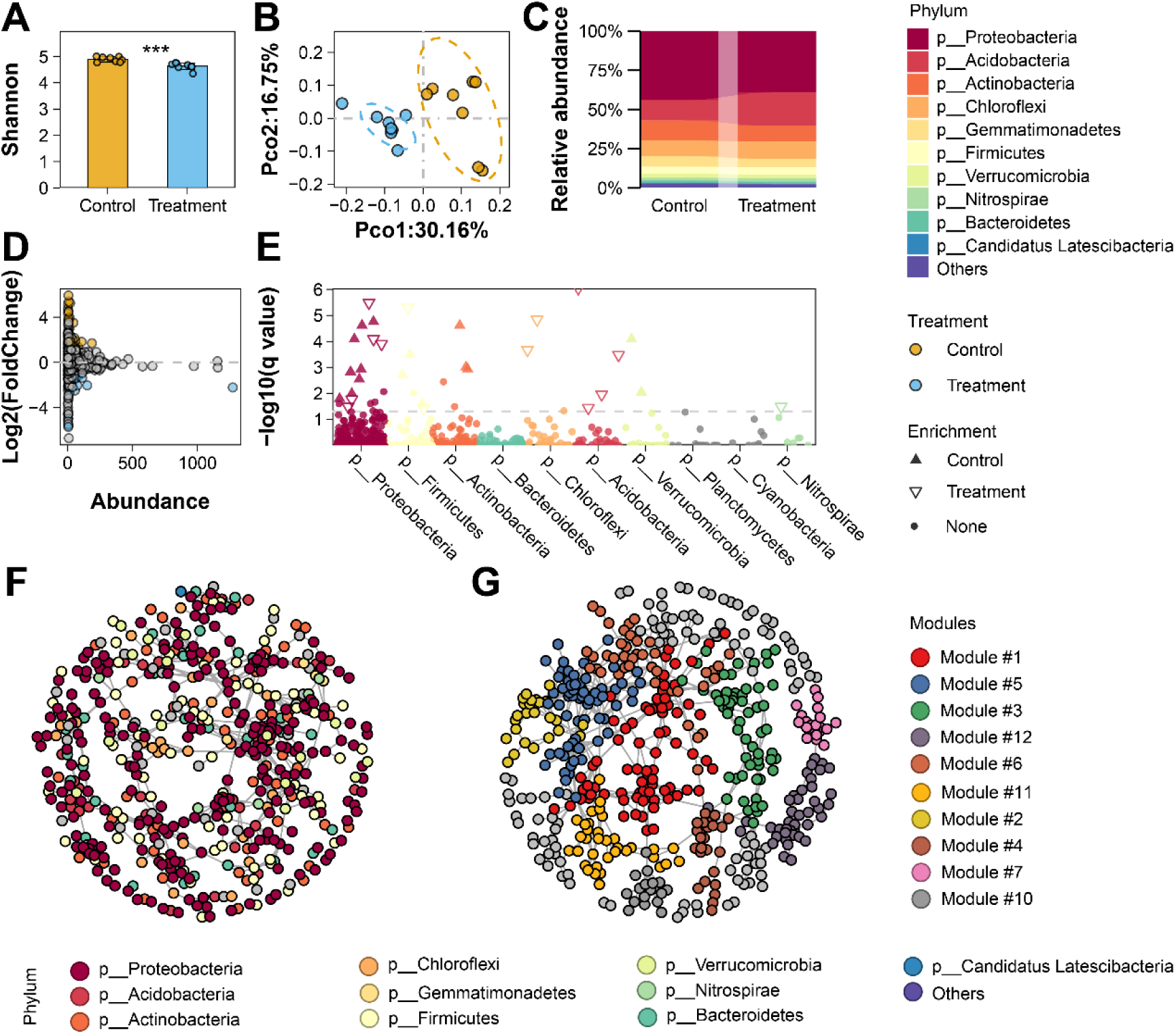
Representative visualizations generated from the sample dataset in the *LorMe* package. (A) Bar plot of the Shannon diversity index, with statistical differences indicated by t-tests (***: p < 0.001). (B) PCoA ordination illustrating differences in community structure; ellipses represent 85% confidence intervals. (C) Stacked bar chart showing the phylum-level composition of the ten most abundant phyla. (D) Volcano plot displaying taxon abundance versus log₂ fold change based on DESeq2, with colors indicating significantly differential taxa. (E) Manhattan-style plot showing phylum-level taxonomy and log₂ fold change, with points colored by phylum. (F-G) Co-occurrence networks highlighting (F) phylum-level taxonomy and (G) module structure.

When applied to the demonstration dataset containing sixteen samples with a two-group design and approximately twenty-two thousand ASVs, the pipeline completed the full analysis suite, including alpha diversity profiling with six indices and three visualization styles, beta diversity ordination using three methods with corresponding visual outputs, differential taxa identification using three statistical approaches with four visualization styles, and co-occurrence network construction for each treatment as well as the combined dataset. The entire workflow executed within fifteen seconds (user time 0.17 s, system time 0.04 s, elapsed time 14.77 s) on a laptop with an Intel i7 processor and 16 GB RAM. The pipeline automatically generated a structured results directory containing thirty-seven high-resolution figures, eighteen tidy data tables, five serialized analysis objects, and a comprehensive analysis log documenting the workflow steps, method selection, parameter settings, and any skipped or failed operations. All visual outputs were accompanied by source data and re-visualization functions, allowing flexible downstream exploration and customization. To balance convenience and analytical flexibility, a total of forty-four adjustable parameters are embedded within the package environment with predefined defaults. The pipeline function remains concise by internally invoking these options, while users maintain the ability to modify settings when deeper exploration is required (Supplementary Table 2). By default, the pipeline performs the entire workflow, although specific modules can be selectively executed as needed. Collectively, this demonstration highlights the design philosophy of the *LorMe* package, emphasizing lightweight architecture, ease of use, low entry barrier, and high reproducibility. Analytical tasks that traditionally require extensive scripting, numerous function calls, and manual parameter tuning can now be completed with minimal user input, making advanced microbial ecological analyses accessible to non-expert users without compromising methodological integrity.

### 3.3 Comparison with existing Tools

We compared *LorMe* with three widely used R packages for microbiome data analysis: *phyloseq*, *microeco*, and MicrobiotaProcess. The comparison focused on modularity, pipeline availability, ecosystem interoperability, supported data types, and output reusability (Table 1). All packages incorporate a modular architecture, but only *LorMe* provides true modular analysis, allowing users to execute individual analytical components, including community feature analysis, differential abundance, and network inference, as standalone modules. Regarding workflow automation, *LorMe* uniquely supports a fully integrated one-command pipeline, whereas *phyloseq* does not include workflow automation and *microeco* and MicrobiotaProcess offer only partial workflow capabilities. *LorMe* further provides full bidirectional interoperability with both *phyloseq* and *microeco*, enabling seamless integration into the broader microbiome R ecosystem. In contrast, *microeco* restricts analyses to its own object format, and MicrobiotaProcess is dependent on *phyloseq* interoperability. All four packages support amplicon and metagenomic data inputs. Similar to the flexible output design of MicrobiotaProcess, *LorMe* offers comprehensive source-data accessibility and downstream export functions, including network tables compatible with visualization platforms such as Gephi and Cytoscape. Collectively, these comparisons highlight the lightweight, flexible, and reproducible design of *LorMe*, which lowers the coding barrier while maintaining broad compatibility and analytical depth across microbiome research workflows.

**Table 1.** Comparison of *LorMe* with three widely used microbiome analysis R packages.

| Package | *Phyloseq*<br>(v1.48.0) | *microeco*<br>(v1.16.0) | MicrobiotaProcess<br>(v1.23.0) | *LorMe*<br>(v2.0.1) |
| --- | --- | --- | --- | --- |
| Platform | Github | CRAN & Github | Github | CRAN & Github |
| Modular design | Yes | Yes | Yes | Yes |
| Modular analysis | No | No | No | Yes |
| Pipeline support | No | Semi-workflow | Semi-workflow | Fully support |
| Interoperability | Limited<br>(*phyloseq* only) | Limited<br>(*microeco* only) | Limited<br>(*phyloseq* only) | Full bidirectional<br>(*phyloseq/microeco*) |
| Data types | Amplicon and metagenomic profiles | Amplicon and metagenomic profiles | Amplicon and metagenomic profiles | Amplicon and metagenomic profiles |
| Output Reusability | Limited | Limited | Flexible | Flexible |

### 4.4 Real Case Reproduction

To demonstrate the practical utility, convenience, and reproducibility of *LorMe* in real-world research scenarios, we applied the complete workflow to an independent dataset profiling the rhizosphere bacterial communities of eggplants with contrasting resistance to bacterial wilt [19]. The case study encompassed all analytical steps, beginning with data loading, followed by encapsulation into a *LorMe* object, global configuration adjustments, execution of the full pipeline, and downstream visualization. Using a single one-command pipeline command, *LorMe* generated a comprehensive suite of analytical outputs, including community diversity metrics, beta-diversity ordination, differential taxa identification, co-occurrence network patterns, and a meta-network highlighting phenotype-sensitive microbial group. The entire workflow required only 36 valid commands and completed within 20 seconds on a standard workstation, demonstrating both the efficiency and user-friendliness of the package. All statistical results, source data, figures, and intermediate objects were automatically archived into a structured results assignment, enabling full reproducibility and convenient downstream customization.

*LorMe* successfully reproduced the key bacterial community features distinguishing resistant and susceptible eggplants (Figure 4). Alpha diversity metrics indicated no significant differences between phenotypes (Figure 4A), whereas PCoA and PERMANOVA revealed clear separation in community structure (Figure 4B). Community composition at the phylum level showed that the ten most abundant phyla accounted for 92.82 percent to 98.10 percent of the total community proportion (Figure 4C). Indicator analysis identified 131 and 110 genera significantly enriched in resistant and susceptible phenotypes, respectively (Figure 4D). The circular Manhattan-style plot highlighted distinct taxonomic enrichment patterns, with resistant plants enriched in Actinobacteria, Firmicutes, and Chloroflexi, while susceptible plants showed enrichment of Bacteroidetes and Verrucomicrobia (Figure 4E). Co-occurrence network analyses indicated comparable network complexity between phenotypes (Figure 4F). The meta-network, integrating co-occurrence structure with indicator taxa, revealed phenotype-associated clustering: resistant-enriched genera were primarily grouped in module 3, whereas susceptible-enriched genera were concentrated in module 2 (Figure 4G). Module-level abundance patterns further confirmed these trends, with module 3 displaying significantly higher relative abundance in the resistant phenotype (p<0.001, t-test), comprising genera from multiple phyla. Module 2, characterized mainly by Proteobacteria and Acidobacteria, exhibited higher abundance in the susceptible phenotype (Figure 4H).

**Figure 4.**
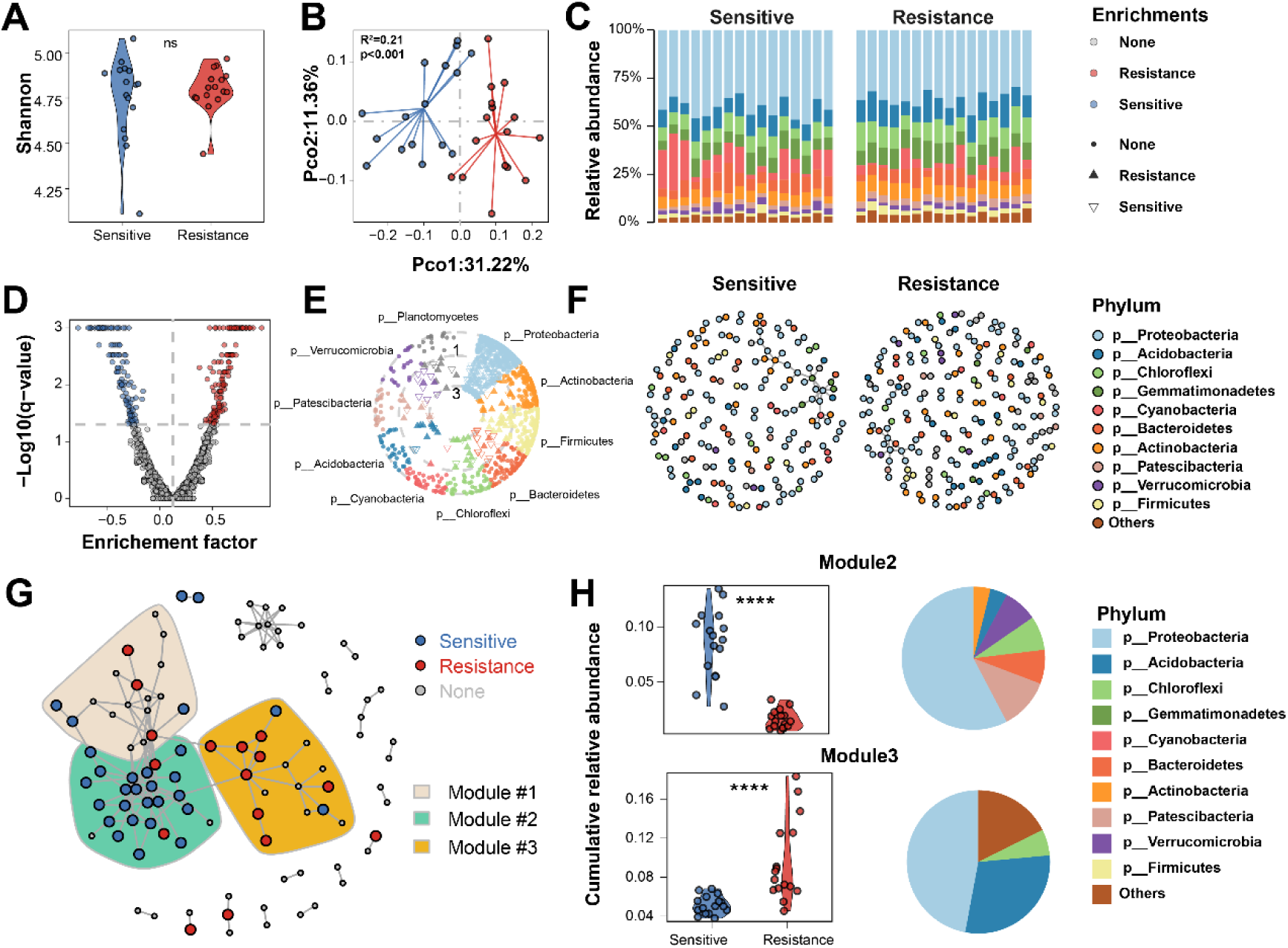
Bacterial community profiles, differential taxa, co-occurrence networks, and meta-network patterns in the real case study analyzed using the *LorMe* package. (A) Violin plot showing bacterial Shannon diversity in the rhizosphere of resistant and sensitive phenotypes. (B) PCoA ordination illustrating differences in bacterial community structure, with PERMANOVA statistics labeled. (C) Stacked bar chart showing the phylum-level composition of the ten most abundant phyla across samples in the two phenotypes. (D) Volcano plot displaying enrichment factor versus log₂ fold change based on indicator analysis, with colors marking significantly differential taxa. (E) Circular Manhattan-style plot presenting phylum-level taxonomy and log₂ fold change, with points colored by phylum. (F) Co-occurrence network constructed using Spearman’s rho ≥ 0.75 and adjusted p < 0.05, with node colors indicating phylum identity. (G) Meta-network integrating indicator taxa and co-occurrence relationships. The three modules containing the largest number of differential taxa are highlighted; node colors indicate whether a taxon is identified as differential. (H) Module abundance and taxonomic composition for the highlighted modules. Asterisks indicate statistical significance based on t-tests (****: p < 0.0001).

Collectively, this real-case application demonstrates capability of *LorMe* to deliver reliable, publication-ready microbiome analyses. The unified encapsulation system, global configuration framework, optimized pipeline architecture, and modular analytical design enable *LorMe* to function as both a comprehensive end-to-end solution and an interoperable component within broader microbiome research workflows.

## Discussion

In this study, we developed *LorMe* (Lightweight One-line Resolving Microbial Ecology), a R package that provides a standardized microbiome analysis framework streamlining the entire analytical lifecycle from raw-data preprocessing to publication-ready visualization. By integrating data encapsulation and unified configuration management within a controllable architecture, *LorMe* effectively addresses long-standing challenges in microbiome research, including fragmented workflows, inconsistent parameter settings, and limited reproducibility. Moreover, *LorMe* achieves bidirectional interoperability with major R-based microbiome analysis ecosystems, enabling seamless data exchange and flexible incorporation of external analytical tools. The package consolidates widely used ecological and statistical analyses into a standardized command system, while further enhancing flexibility through archiving of intermediate objects, source data, and graphical outputs. Notably, the implementation of highly concise one-command pipeline commands substantially lowers the technical threshold for users without compromising analytical rigor or scalability. Collectively, these features position *LorMe* as a robust, user-friendly, and interoperable R package capable of supporting both routine microbiome profiling and advanced multi-layer community analyses.

The comparative evaluation of existing R-based microbiome tools in this study highlights the structural fragmentation that continues to characterize the current analytical ecosystem [6]. Our results demonstrate that *LorMe* effectively alleviates several of these long-standing limitations by providing a unified architecture that bridges previously isolated workflows. Rather than replicating the functionalities of established packages, *LorMe* operates as an interoperable ecological connector that links widely used infrastructures such as *phyloseq* and *microeco* through fully bidirectional object compatibility. This design enables users to integrate analytical functions across ecosystems without reconstructing data objects or restructuring workflows, thereby reducing the cumulative scripting burden and minimizing error propagation across analytical steps. Moreover, the unified configuration management implemented in *LorMe* ensures that analytical parameters remain consistent across modules, addressing a common source of irreproducibility in user-assembled workflows. The results of the one-command pipeline further show that automated workflow standardization does not compromise analytical flexibility; instead, it produces internally coherent outputs that can be directly reused across downstream visualization, interpretation, and reporting. These features collectively position *LorMe* as a methodological framework that enhances interoperability, reduces procedural complexity, and reinforces reproducibility within microbiome research, complementing rather than replacing existing analytical tools.

The practical application of *LorMe* in real research scenarios further demonstrates its ability to generate biologically interpretable and methodologically consistent results. Across its integrated modules, *LorMe* produced coherent patterns that consistently distinguished microbial community responses between experimental treatments. This multi-angle analytical design not only provides complementary views of the same ecological signal but also enables users to examine the robustness of findings across independent methods. For example, identified differential taxa aligned closely with distinct network modules and module-level abundance shifts, echoing commonly reported applications of meta-network frameworks in identifying key microbial responders [20] to environmental or experimental gradients [21]. To support this consistency, *LorMe* retains full transparency and modifiability in each analytic step, allowing users to trace, adjust, or extend any component of the workflow when validating results or integrating additional ecological traits. Taken together, these case-study demonstrations show that *LorMe* not only streamlines microbiome analysis but also provides a reliable and integrative framework for generating insights that can help exploring deeper ecological mechanisms.

Despite its comprehensive capabilities, *LorMe* has several limitations that should be acknowledged. First, although the pipeline supports both amplicon and metagenomic microbial profile data, current implementations offer limited functionality for phylogeny-informed analyses, such as β NTI calculation for community assembly processes and phylogeny-aware analysis [22]. Second, analyses at the functional gene level, including gene-microbe correlations or functional enrichment, are not yet integrated into the main workflow. Third, while *LorMe* provides multiple visualization templates, the range of graphic styles and themes remains relatively limited compared with fully customizable plotting frameworks. These limitations do not compromise core analyses but represent opportunities for future improvement. Looking forward, *LorMe* is poised for continued development. Planned extensions include 1) the integration or interoperation of new analytical methods, such as MBPD [23] and specialized tools for network analysis [24]. 2) Expansion of visualization templates. 3) Development of a user-friendly Shiny application to facilitate interactive exploration [25]. Together, these developments position *LorMe* as a flexible, interoperable, and extensible framework that can continue to evolve alongside the rapidly advancing microbiome research landscape.

## Availability and requirements

Project name: LorMe (Lightweight One-Line Resolving Microbial Ecology)

Project home page: https://github.com/wangnq111/LorMe

Official/formal distribution: https://cran.r-project.org/web/packages/LorMe/index.html

Help documentation: https://wangnq111.github.io/Gettingstarted.html

Operating system(s): Linux, OS X, Windows

Programming language: R

License: GPL-3

## Availability of data and materials

The script of sample and real case demonstration of this article is available on GitHub at https://github.com/wangnq111/LorMe_examples

## Funding

This research was supported by the National Natural Scientific Foundation of China 42325704, the Natural Science Foundation of Jiangsu Province BK20230988 and the National Natural Scientific Foundation of China W2431030. We sincerely thank Ye Tao and Liang Zeng from Shanghai Biozeron biotechnology Co. LTD for the support in designing the package. We appreciate users’ feedback in improving functions of package in beta version.

## Author information

Ningqi Wang and Yaozhong Zhang have equal contributions.

## Contributions

Ningqi Wang: Conceptualization, Writing – original draft, Writing – review & editing, Visualization, Formal analysis.

Yaozhong Zhang: Visualization, Data curation, Formal analysis, Writing – review & editing. Xinrun Yang: Investigation

Yang Gao: Funding acquisition

Hang Xia: Data curation.

Yangchun Xu: Supervision.

Qirong Shen: Supervision.

Alexandre Jousset: Conceptualization.

Gaofei Jiang: Conceptualization, Investigation, Writing – review & editing.

Zhong Wei: Conceptualization, Investigation, Writing – review & editing, Funding acquisition.

## Ethics declarations

### Ethics approval and consent to participate

Not applicable.

### Consent for publication

Not applicable.

### Competing interests

The authors declare that they have no competing interests.

## Supplementary Information

Supplementary Tables contain:

Supplementary Table 1: Detailed list of outputs in LorMe pipeline analysis.

Supplementary Table 2: Detailed list of parameters in LorMe pipeline analysis.

## Notes

https://github.com/wangnq111/LorMe_examples

### Competing Interest Statement

The authors have declared no competing interest.

